# Probabilistic Cardiac Digital Twins for Robust Patient-Specific Modeling

**DOI:** 10.64898/2026.05.07.723610

**Authors:** Dimitris G Giovanis, Kelly Zhang, Justin Tso, Mauro Maggioni, Ioannis G Kevrekidis, Natalia Trayanova

## Abstract

Uncertainty quantification (UQ) in computational heart models is essential for reliable cardiac digital twins (DTs) in personalized medicine, yet remains challenging. Traditional Monte Carlo and stochastic Galerkin methods often become impractical in the high-dimensional, nonlinear state variable and parameter spaces of cardiac electrophysiology and mechanics. This article introduces a framework for learning a joint probability density over cardiac observables and model parameters, enabling the characterization of statistical dependencies across a large number of variables in patient-specific cardiac DTs. By sampling from this density and conditioning on available data, useful predictive distributions can be constructed, allowing uncertainty to be propagated through the model and quantified in terms of variability. Conditional regression can then be performed directly on this learned density, enabling systematic exploration of interdependencies among observables for both predictive inference and model design. The statistical methodology adopts a geometry-aware generative learning framework, recently introduced by the authors, that decouples the learning of data geometry from sampling. First it identifies a low-dimensional latent representation that captures the intrinsic structure of the data and its multiscale geometric features. A stochastic differential equation is then formulated directly in the low-dimensional latent space to generate samples efficiently; these are subsequently mapped back to the high-dimensional space of cardiac states and parameters through a smooth lifting operator. We demonstrate the approach on a ventricular arrhythmia prediction benchmark, where the learned joint probability density enables the construction of predictive distributions over key parameters (e.g., conductivities, fibrosis patterns) through sampling and conditioning. This enables uncertainty to be propagated and quantified through sampling and conditioning on the learned joint density, with substantially fewer model evaluations than conventional UQ methods.

## Introduction

Computational modeling of the human heart has become a cornerstone of modern cardiovascular research, enabling *in-silico* investigations of electrophysiological (EP) mechanisms, disease progression, and treatment outcomes. Recent advances in cardiac CT have allowed the integration of patient-specific anatomy and tissue remodeling distributions for individualized predictions that can help diagnoses, therapy planning, and risk stratification. DTs have been used in several studies of heart rhythm dysfunction, both atrial and ventricular. They have demonstrated accurate predictions of the risk of sudden cardiac death (1) by evaluating the arrhythmogenic propensity of the disease-remodeled myocardium. Cardiac DTs that incorporate genetic predisposition (2) or penetrating adiposity (3) have recently been used to predict the locations of ventricular tachycardias (VTs), lethal rhythm disorders. DTs have been used to predict noninvasively, before the procedure, the optimal personalized targets in patients undergoing ablation (4) guiding the delivery of therapeutic lesions. However, despite these advances, a persistent challenge remains: performing robust uncertainty quantification (UQ) across the multiple scales (i.e. from cell to organ) that govern cardiac electrophysiology (EP) behavior. In DT predicitons, notable sources of uncertainty are the parameters describing ion channel conductances, gating kinetics, and tissue conductivities. These values range greatly between individuals and cannot be easily measured (5). Consequently, current DTs often rely on population-averaged values or limited experimental datasets, which may not be representative of the true parameter distributions. Understanding how these uncertainties in model input affect simulation-based predictions of arrhythmia inducibility and its dynamics is essential for strengthening the clinical reliability of cardiac DTs.

Current UQ methods typically rely on Monte Carlo sampling, polynomial chaos expansions, or stochastic finite element frameworks (6, 7). However, these frameworks are limited in the context of cardiac DTs by the need to repeatedly evaluate an already expensive forward model. Specifically, these DTs typically couple sets of ordinary differential equations describing action potential behavior to partial differential equations governing electrical wave propagation. These equations are solved over detailed heart geometries represented by 3D computational meshes that can contain up to millions of nodes. Hence, even a few seconds of simulated activity can require hours of computation. Classical UQ using the aforementioned approaches requires an extensive sample of such simulations to explore variability across high-dimensional parameter spaces involving ionic kinetics, conduction properties and complex anatomical features. Insufficient sampling can lead to noisy or unstable estimates of uncertainty, while adequately capturing nonlinear interactions and clinically important events like VT demands prohibitively large numbers of samples. Surrogate-based methods like Gaussian processes (8) or reduced-order models (9, 10) can approximate uncertainty propagation at lower computational cost (11). However, they often fail to capture additional sources of variability such as EP heterogeneity (e.g. repolarization and anisotropy differences) and structural remodeling distributions (e.g. regions of fibrosis or dense scar) which can greatly impact conduction properties. Consequently, achieving an efficient, data-driven UQ that preserves both computational tractability and physiological consistency remains an open challenge in the development of cardiac DT.

In this context, recent advances in generative modeling offer a promising alternative by enabling the direct learning of high-dimensional probability distributions from limited data, thereby reducing reliance on repeated forward simulations. Methods such as variational autoencoders (VAEs) (12, 13), normalizing flows (14), implicit models (e.g., generative adversarial networks (15, 16)), and score-based diffusion models (17–19) have demonstrated exceptional capabilities in sampling from complex distributions without requiring explicit likelihood formulations. These methods have demonstrated state-of-the-art performance in image (20), audio (21), graph generation (22), and protein chemistry (23), among others. Despite their empirical success, these models face several limitations when applied to scientific datasets. Specifically, they often rely on unconstrained Euclidean embeddings, which can obscure the low-dimensional, nonlinear structures, such as anatomical surfaces and biophysical parameter spaces. As such, these models may require large amounts of data or extensive training to accurately learn the geometry of the underlying distribution.

To address these limitations, generative modeling can be combined with manifold learning, which leverages the observation that high-dimensional scientific data often lie on low-dimensional, nonlinear manifolds embedded in ambient space (24). Examples of manifold learning methods include Isomap (25), local linear embedding (LLE) (26), and Diffusion Maps (27). In this context, Probabilistic Learning on Manifolds (PLoM) (28–31) approximates a data distribution by combining kernel density estimation, Diffusion Maps (DM), and an Itô stochastic differential equation (ISDE) to generate samples constrained to a learned manifold (28–31). Building on these ideas, we introduce a data-driven framework that learns a joint probability density over cardiac observables and model parameters. This representation enables uncertainty propagation through sampling, patient-specific inference via conditioning on available data, and systematic analysis of dependencies through conditional regression. By operating on intrinsic low-dimensional manifolds, the approach overcomes the computational limitations of classical UQ methods while preserving physiological consistency.

To realize this framework in the context of cardiac digital twins, we adopt a geometry-aware generative approach based on Double Diffusion Maps Probabilistic Learning on Manifolds (DD-PLoM) (32). DD-PLoM extends manifold-based probabilistic learning by enabling efficient sampling directly in a learned latent space, facilitating exploration of complex, non-Euclidean parameter distributions without requiring explicit likelihoods or intrusive solver modifications. When coupled with a 3D DT of the electrical behavior of a patient’s heart, this approach generates physiologically consistent realizations of wavefront dynamics under electrophysiological parameter uncertainty, supporting robust prediction of arrhythmia risk.

Within this setting, we build upon recent advances in high-fidelity cardiac modeling [REF] by integrating cellular electrophysiology and tissue-level conduction variability into a unified, patient-specific biventricular reconstruction that serves as a foundation for uncertainty-aware simulations. Our contributions are threefold:

1. We introduce a framework for learning joint probability densities over cardiac observables and model parameters, enabling uncertainty propagation, conditioning, and dependency analysis within patient-specific cardiac DTs.
2. We deploy a geometry-aware generative approach (DD-PLoM) that operates on intrinsic manifolds, enabling efficient sampling of physiologically consistent parameter distributions without intrusive solver modifications.
3. We show that the proposed approach efficiently reconstructs posterior distributions of key cardiac features using significantly fewer samples than conventional UQ techniques, thereby enabling real-time, generative learning for personalized cardiac models.

In the remainder of this paper, we present a clinically relevant application of the proposed framework in a cardiac digital twin of a patient with repaired Tetralogy of Fallot (rToF) experiencing ventricular tachycardia (VT). Section reports validation results and demonstrates how the framework distinguishes between arrhythmic and non-arrhythmic simulations through conditional inference. Section A describes the methodology, Section D interprets the findings in a clinical context, and Section I summarizes the main contributions and outlines directions for clinical translation.

## Results

Tetralogy of Fallot (ToF) is the most common form of complex congenital heart disease (33). While surgical repair at a young age is highly successful, surgical scar and implanted material create anatomical isthmuses that may harbor VT circuits for adult patients. These isthmuses can be detected using noninvasive methods such as late gadolinium enhancement cardiac magnetic resonance imaging (LGE-CMR) which visualizes structural changes in the heart (scar/fibrosis or surgical repair material). However, not all rToF patients with isthmuses are inducible for VT, suggesting that uncertainties in conduction velocity and action potential dynamics play an important role. To enable data-driven learning of the joint distribution over cardiac observables and electrophysiological parameters, a patient-specific biventricular model of a rTOF heart was constructed from 3D LGE-CMR data, and VT induction was simulated under 100 distinct EP parameter sets.

Across the 100 simulations, the VT induction protocol produced a wide range of outcomes. Thirty–nine parameter sets resulted in sustained activity (lasting ≥2 s), and sixty simulations of the patient’s DT resulted in non-sustained arrhythmic activity or were non–inducible for ventricular arrhythmia. One parameter set was excluded because the paced wavefront failed to propagate with the prescribed stimulus strength. Throughout this section, we use the term “VT” to describe only sustained activity, and “non–VT” for both un-sustained activity and non-inducibility. VT was consistently induced using a rapid pacing protocol, by either the second or third premature stimulus (S2–S3), with an average arrhythmia cycle length of 286±5.4 ms. Representative reentrant patterns are shown in Fig. 1; each row illustrates snapshots of membrane voltage over time for two example VT circuits induced under distinct parameter sets.

**Fig. 1.**
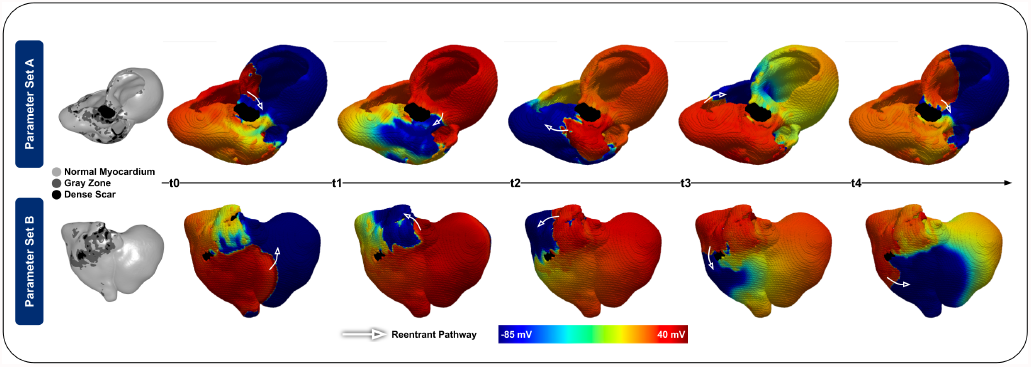
Representative VT morphologies.

### Parameter and response variability in VT vs. non–VT DTs

We first examined how variability in model parameters and EP responses differed between DT simulations that resulted in VT induction and those that did not. Figure 2 summarizes the distributions of all sampled parameters. We estimate uncertainty in the kernel density estimate (KDE) using nonparametric bootstrap resampling. Specifically, we repeatedly resample the data with replacement, compute a KDE for each resample, and construct pointwise standard deviation bands from the empirical variance of the bootstrapped density evaluations.

**Fig. 2.**
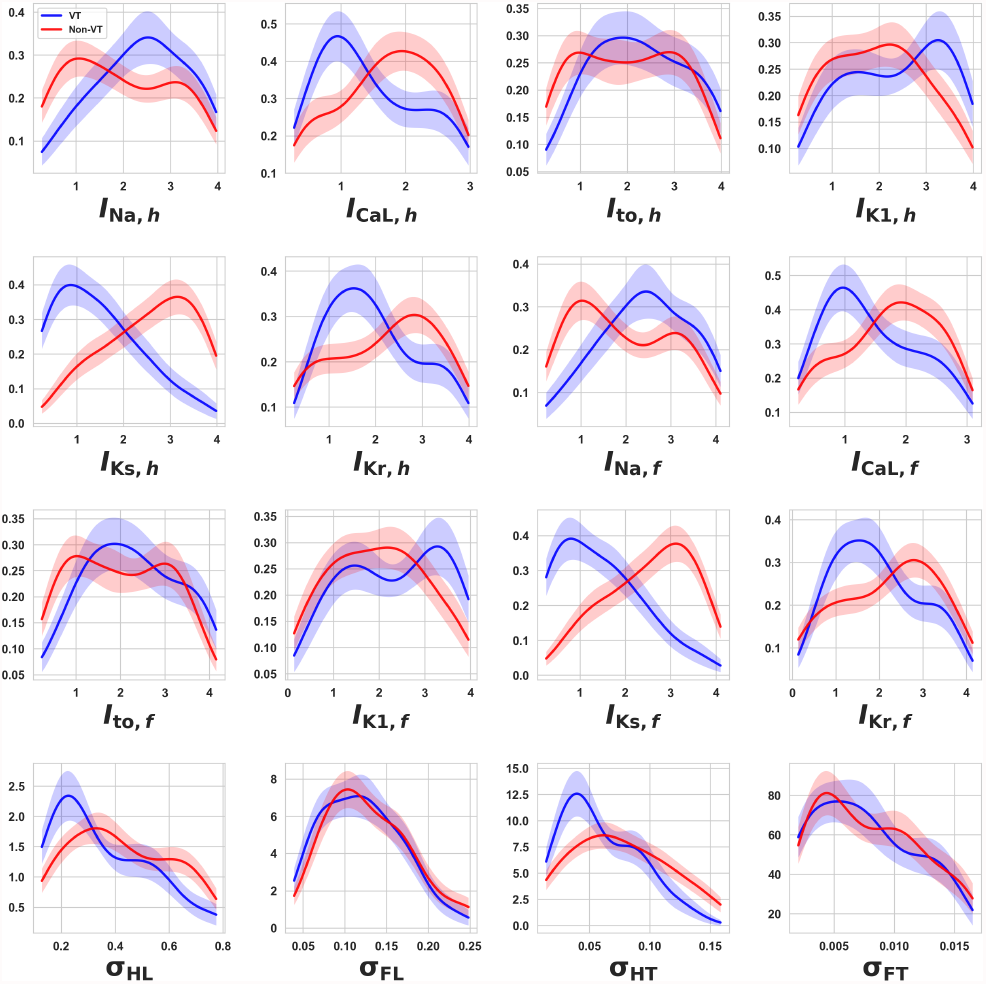
Marginal distributions of cell-level EP and tissue properties for VT (arrhythmia) and non-VT (non-arrhythmia) simulations. We estimate uncertainty in the kernel density estimates using nonparametric bootstrap resampling. Specifically, we repeatedly resample the data with replacement, compute a KDE for each resample, and construct pointwise standard deviation bands from the empirical variance of the bootstrapped density evaluations.

At the cell level, the sampled conductance scalings cover broad but physiologically plausible ranges for fast sodium (INa), L–type calcium (ICaL), inward rectifier potassium (IK1), transient outward potassium (Ito), and the slow/fast delayed rectifier potassium currents (IKs, IKr), consistent with the design of the cell-population modeling pipeline. Several parameters have pronounced separations between the VT and non-VT marginal distributions, which indicate that variations in these parameters individually exert larger influence on arrhythmogenic propensity within the sampled parameter space. Among VT-inducing simulations, INa was likely to be upregulated, consistent with the expected effect of augmented sodium conductance in enhancing excitability and lowering the threshold for sustained propagation. Higher VT likelihood was also associated with reduced ICaL, a trend that aligns with shorter AP durations and refractory periods which can facilitate reentry. A similar pattern was observed for IKs, where lower conductance values tended toward greater VT incidence. This shift may reflect diminished repolarization reserve and increased susceptibility to conduction block between healthy and fibrotic tissue. Together, these marginal distributions identify INa, ICaL, and IKs as primary parameters that, in isolation, could bias the system toward arrhythmogenic behavior.

At the tissue scale, the simulated population also shows substantial variability in conduction velocity (CV) and anisotropy across healthy and diffuse fibrosis regions. By construction, healthy longitudinal CV (*CV*_*HL*_) ranges from 60–130 cm/s and fibrotic transverse CV (*CV*_*FT*_) from 18–30 cm/s. Within these ranges, VT-inducible simulations tend to occupy the lower end of healthy longitudinal and transverse conductivities (*σ*_*HL*_, *σ*_*HT*_). In contrast, conductivities of fibrotic regions (*σ*_*FL*_, *σ*_*FT*_) show less obvious separation between the two groups. While the surgical patch and associated fibrotic isthmuses define potential reentrant pathways, slowed conduction in surrounding healthy myocardium shortens the circuit wavelength and allows sufficient recovery time for re-excitation of the myocardium, enabling sustained VT. This observed association between reduced conduction velocity in healthy tissue and simulations resulting in VT is consistent with established mechanisms of reentry in the disease.

Figure 3 shows transmembrane potentials measured from a point within a clinically important isthmus bounded by the surgical patch and pulmonary valve annulus. For each group (VT on the left in blue, non–VT on the right in red), the dashed line denotes the mean trace and the shaded region represents one standard deviation across all simulations in that group. The initial depolarization peaks in the averaged traces, between 0 ms to 500 ms, correspond to the final beats of a rapid pacing train. In simulations that resulted in VT, arrhythmia was typically initiated by the second or third premature stimulus; accordingly, the oscillatory activity observed between approximately 500 ms and 1000 ms reflects wavefronts elicited by these premature beats. Following this interval, the emergence of more regular, periodic depolarizations corresponds to self-sustaining VT wavefronts. In contrast, non-VT simulations exhibit longer oscillations during the premature stimulation phase, as premature beats were delivered up to four times. Because no self-sustaining activity developed in these simulations, electrical activity ceased thereafter, and the remaining portion of the traces was padded with zeros, resulting in a flat signal.

**Fig. 3.**
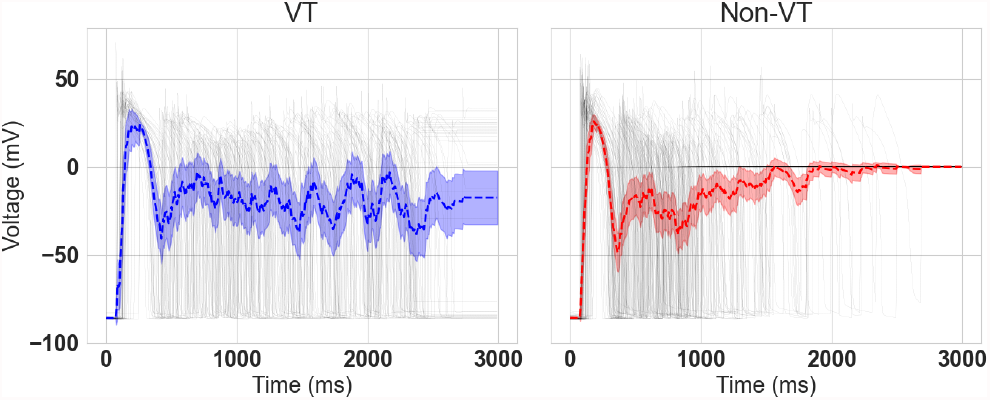
Mean trans-membrane voltage time courses (dashed lines) and variability bands (shaded regions, one standard deviation) for simulations that developed VT (left, blue) and those that remained non-inducible (right, red). With grey we see the individual trans-membrane voltage time courses.

### A. Augmentation of the learned dataset

We enrich the limited dataset by generating a large ensemble of synthetic samples from the learned joint distribution *p*(**W, R**). In our setting, **W** ∈ ℝ^16^ denotes the vector of input parameters (cell-level and tissue-level properties) and **R** ∈ ℝ^582^ represents the corresponding quantities of interest. Starting from the original dataset of 100 samples, we first perform a clustering analysis (k-means) to separate VT from non-VT cases, yielding a VT dataset with 39 VT samples and a non-VT dataset with 61 samples. We emphasize that sampling such a distribution is particularly challenging due to the combination of high dimensionality and limited data. With only 39 and 61 samples in a 16+582 = 596-dimensional space, the observations occupy an extremely sparse region, leaving most of the space effectively unexplored. As a result, estimating the underlying distribution becomes ill-posed, as many distinct distributions can be consistent with the available data. In addition, the curse of dimensionality renders distances between points less informative, making it difficult to define meaningful neighborhoods or accurately estimate densities. This poses a fundamental challenge for generative modeling, which must infer structure in regions where no data are observed, increasing the risk of generating unrealistic or non-physical samples.

Using the DD-PLoM approach, we generate 100 copies of each dataset. This results in 3,900 synthetic samples for the VT group and 6,100 for the non-inducing group. We note that, since the generative model is trained to learn the joint distribution *p*(**W, R**), it is essential that the augmented marginals remain close to the reference marginals to ensure that this joint structure is accurately captured. However, while agreement at the marginal level is important, our primary interest lies in accurately capturing the conditional relationships encoded in the joint distribution. This becomes particularly important when performing conditional sampling (e.g., sampling **R** |**W**), as any distortion in the marginals of **W** can propagate through the conditioning step and lead to biased or non-physical response predictions.

Figure 4 compares reference trajectories with simulated trajectories generated by the learned distribution for both arrhythmic (VT) and non-arrhythmic regimes. In the VT case, the augmented signals accurately reproduce the irregular oscillatory patterns and amplitude variations observed in the reference data, capturing both the transient behavior and sustained fluctuations characteristic of ventricular tachycardia. Despite the increased complexity of the dynamics, the generated responses remain well-aligned with the reference trajectory, indicating that the model preserves the essential temporal structure of the system. In the non-VT regime, where the dynamics are more regular and gradually damped, the agreement between reference and augmented signals is even tighter, with minimal deviations over time. This reflects the reduced variability and smoother evolution of the underlying system in the absence of arrhythmic activity. Overall, the close correspondence between reference and augmented responses across both regimes demonstrates that the proposed approach effectively captures regime-dependent dynamics while maintaining reliable fidelity in the generated signals.

**Fig. 4.**
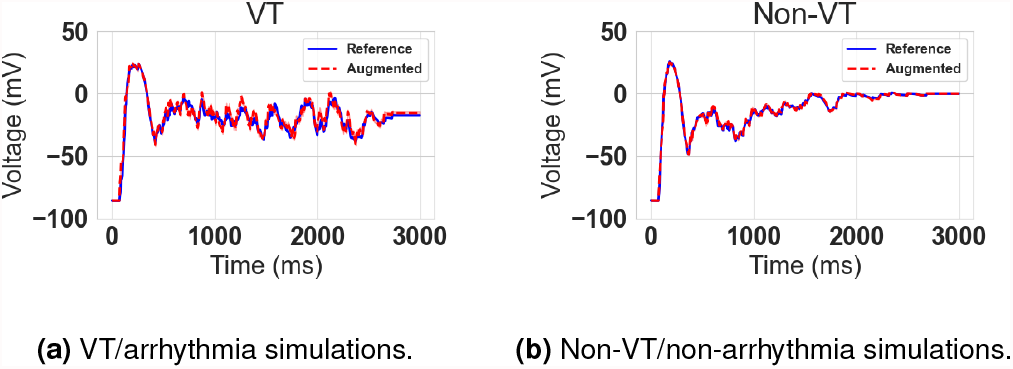
Mean action potential traces obtained with the proposed approach, compared with reference simulations in (a) VT/arrhythmia and (b) non-VT/non-arrhythmia groups.

Figure 5 examines the distributions of transmembrane voltage at four representative time points during the AP upstroke and early repolarization (*t* = 0.07, 0.14, 0.34, and 0.68 ms) for the VT/arrhythmia (left) and non-VT/non-arrhythmia (right) groups. In each panel, the solid curve is the reference kernel density estimate from the 100 simulated DTs, and the dashed curve is the density reconstructed from the learned joint distribution. For both VT and non-VT groups and at all time points, the generative model closely reproduces the location and spread of the voltage distributions. In the arrhythmia group, DD–PLoM *captures the transient broadening and the emergence of bimodal structure during early repolarization*, while in the non-arrhythmia group it *recovers the sharper, highly concentrated peaks* that appear once the wavefront has stabilized. Remaining discrepancies are small and mainly confined to the tails and secondary peaks, indicating that DD–PLoM preserves the essential group-level statistics of the voltage field over time.

**Fig. 5.**
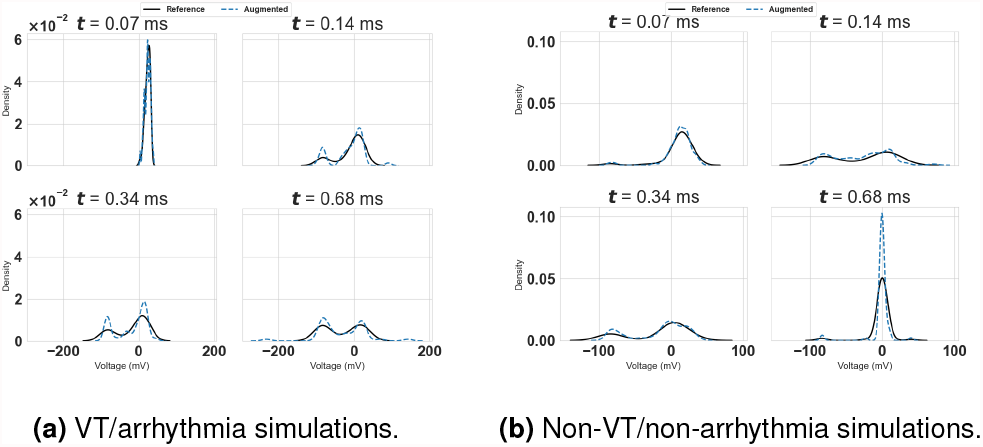
Time-resolved distributions of transmembrane voltage for (a) VT/arrhythmia and (b) non-VT/non-arrhythmia simulations. Solid black curves: reference kernel density estimates from 100 digital-twin simulations. Blue dashed curves: densities obtained from 10,000 DD–PLoM realizations. Snapshots are shown at four representative time points (*t* = 0.07, 0.14, 0.34, and 0.68 ms).

Figures 6(a-b) illustrate the conditional response of the transmembrane potential for four representative parameter configurations **w**_1_, …, **w**_4_, for the VT and the non-VT cases, respectively. Each subplot shows the conditional mean trajectory together with the associated variability, represented by the shaded region corresponding to one standard deviation. Notably, the magnitude of variability is not uniform in time: uncertainty is amplified when the dynamics are highly non-linear. These results demonstrate that the proposed frame-work enables conditional inference of cardiac dynamics that is both physiologically meaningful and uncertainty-aware, providing a data-driven representation of how variability in electrophysiological parameters translates into variability in observable cardiac responses.

**Fig. 6.**
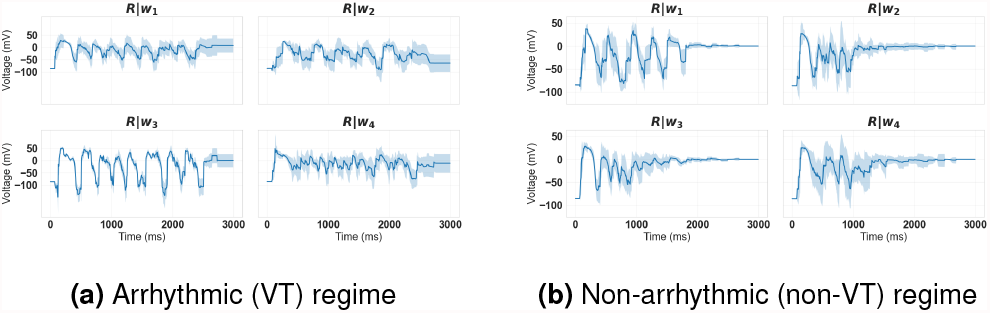
Conditional transmembrane potential trajectories for four representative electrophysiological parameter configurations **w**_1_, …, **w**_4_ across two dynamical regimes. Left: arrhythmic (VT) case. Right: non-arrhythmic case. In each subplot, the solid line denotes the conditional mean, while the shaded region represents one standard deviation derived from the conditional ensemble. The comparison highlights the increased variability and irregular dynamics in the VT regime versus the more stable and damped responses in the non-VT regime.

Figures 7 and 8 demonstrate the ability of the proposed non-parametric conditional sampling framework to generate realistic arrhythmic responses conditioned on specific parameter configurations. Across all plots, the conditional samples (red, dashed) closely follow the reference trajectories (blue, solid), capturing key features such as activation patterns, oscillatory behavior, and repolarization dynamics. Notably, improved agreement is observed for smaller distances *d*, indicating that the local kernel weighting effectively emphasizes samples that are closer in parameter space to the conditioning point. As *d* increases, slight discrepancies emerge, particularly in amplitude and timing, which reflect the increased uncertainty and variability inherent in regions of lower data density. Importantly, even in these cases, the generated samples preserve the qualitative structure of the arrhythmic dynamics, suggesting that the learned joint distribution captures the underlying manifold of physiologically plausible responses.

**Fig. 7.**
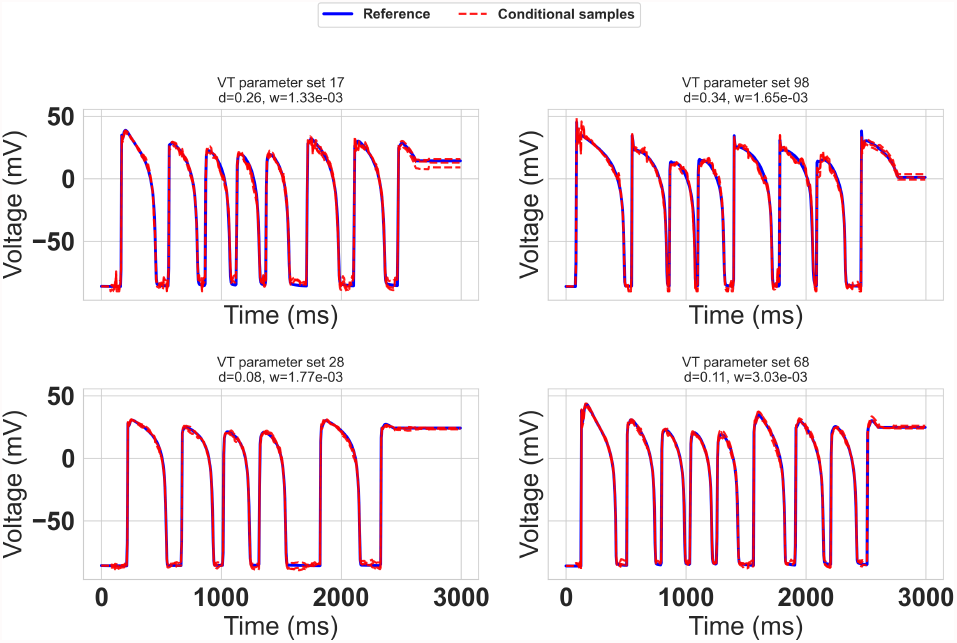
VT case: Comparison between reference transmembrane potential traces (blue, solid) and randomly drawn conditional samples (red, dashed) for multiple VT/arrhythmia-inducing parameter sets.

**Fig. 8.**
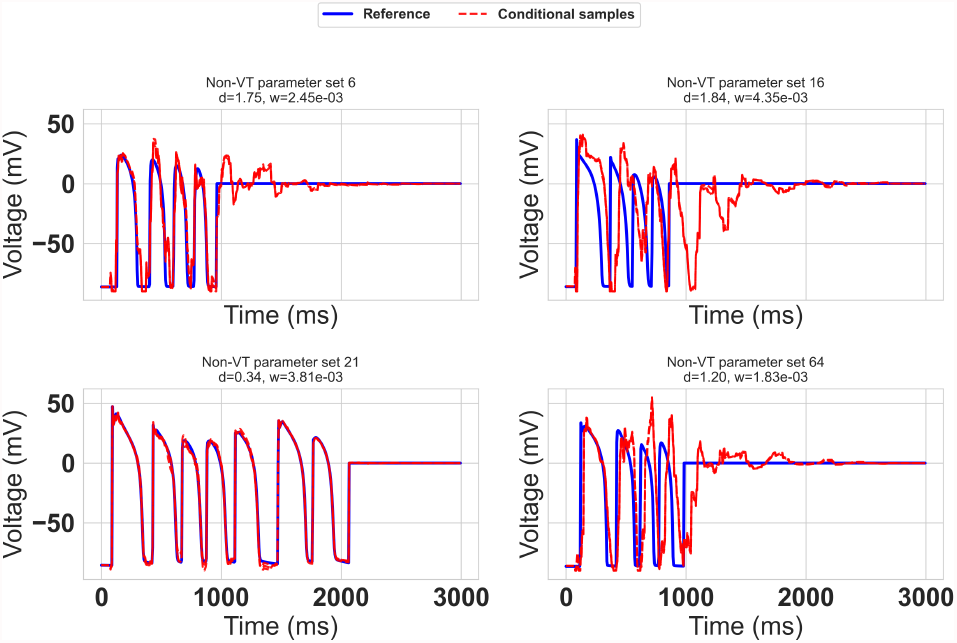
non-VT case: Comparison between reference transmembrane potential traces (blue, solid) and randomly drawn conditional samples (red, dashed) for multiple VT/arrhythmia-inducing parameter sets.

Figures 9 and 10 present conditional sampling results from the learned joint distribution *p*(*R, Q* | *W* = *w*_0_) for two qualitatively distinct physiological regimes: VT and non-arrhythmic dynamics. In both cases, the top-left panels show ensembles of conditional realizations in the observable space *R*, illustrating the variability induced by conditioning on a fixed input *w*_0_. In the VT regime (Fig. 9), the samples exhibit sustained irregular oscillations characteristic of arrhythmic behavior, with significant variability across realizations. In contrast, the non-VT regime (Fig. 10) displays more regular and damped dynamics, reflecting stable cardiac responses with reduced variability. The top-right panels compare the analytical conditional expectation 𝔼[*R* | *W* = *w*_0_] with the empirical mean estimated from conditional samples. In both regimes, the close agreement between these curves demonstrates the accuracy of the conditional expectation estimator and confirms that the sampling procedure is consistent with the learned conditional distribution. The bottom-left panels assess first-order moment consistency in the latent space *Q*. The empirical mean of sampled latent trajectories and the mean obtained from the local Gaussian approximation are both in strong agreement with the model-implied conditional mean. This indicates that the learned representation captures the dominant conditional structure of the data and that both sampling strategies preserve first-order statistics. The bottom-right panels evaluate second-order moment consistency through a comparison of standard deviations. The empirical variability of samples closely follows the theoretical standard deviation derived from the conditional covariance, with the Gaussian approximation providing a reasonable match across most time instances. Notably, in the VT regime, larger deviations and increased variance reflect the inherently more complex and unstable dynamics, while the non-VT case exhibits lower variance and tighter agreement across all estimators.

**Fig. 9.**
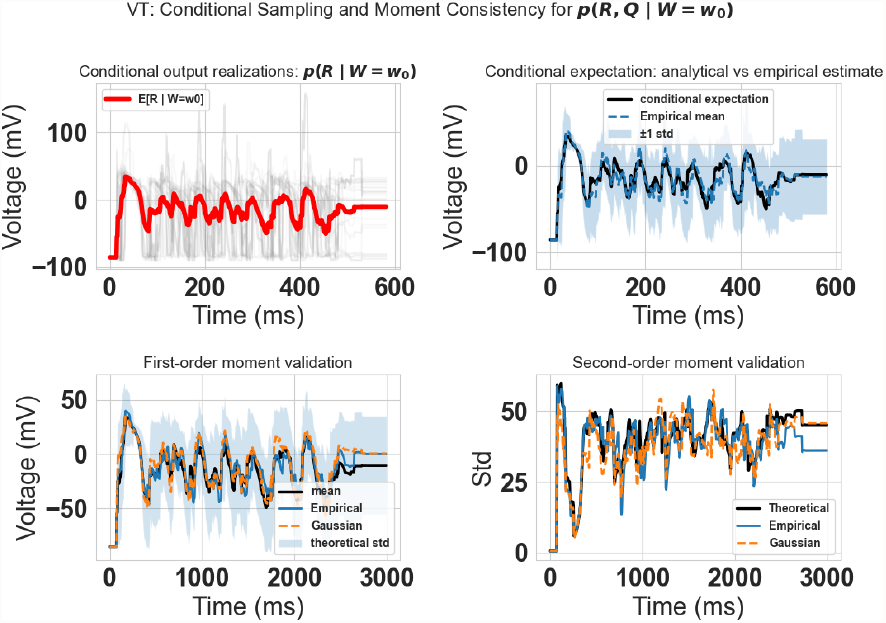
Conditional sampling from the learned distribution *p*(*R, Q* | *W* = *w*_0_) in the arrhythmic (VT) regime. The figure illustrates the variability of conditional realizations and the consistency of first- and second-order statistics in the presence of sustained ventricular activity.

**Fig. 10.**
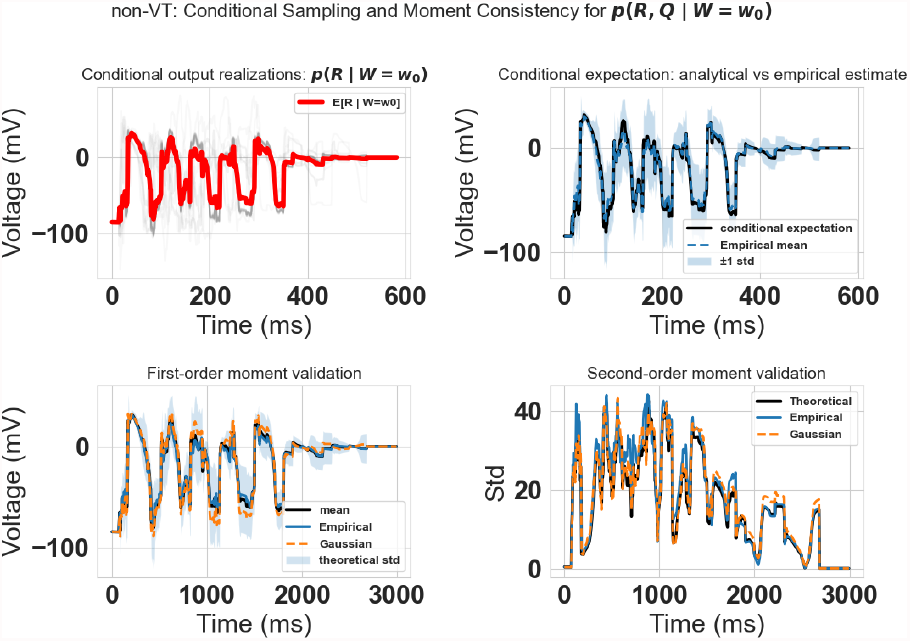
Conditional sampling from the learned distribution *p*(*R, Q* | *W* = *w*_0_) in the non-arrhythmic regime. The model captures stable cardiac dynamics while preserving the statistical structure of the conditional distribution.

Figures 11 and 12 present conditional predictions for unseen validation inputs in the arrhythmic (VT) and non-arrhythmic regimes, respectively. In both cases, the conditional mean predictions closely follow the reference trajectories, demonstrating that the learned model generalizes well beyond the training data. In the VT regime, characterized by sustained and irregular oscillatory activity, the model captures the overall temporal structure and major transitions, while the predictive uncertainty appropriately increases during highly dynamic phases. This reflects the intrinsic variability of the underlying dynamics and the increased difficulty of prediction in this regime. In contrast, the non-VT regime exhibits more regular and damped responses, and the model achieves tighter agreement with the reference signals, accompanied by reduced uncertainty bands. Across both regimes, the reference trajectories are largely contained within the predicted confidence intervals, indicating that the learned conditional distribution provides a reliable quantification of uncertainty. These results highlight the ability of the proposed framework to produce accurate and statistically consistent conditional predictions, while adapting to markedly different dynamical behaviors in both stable and highly nonlinear settings.

**Fig. 11.**
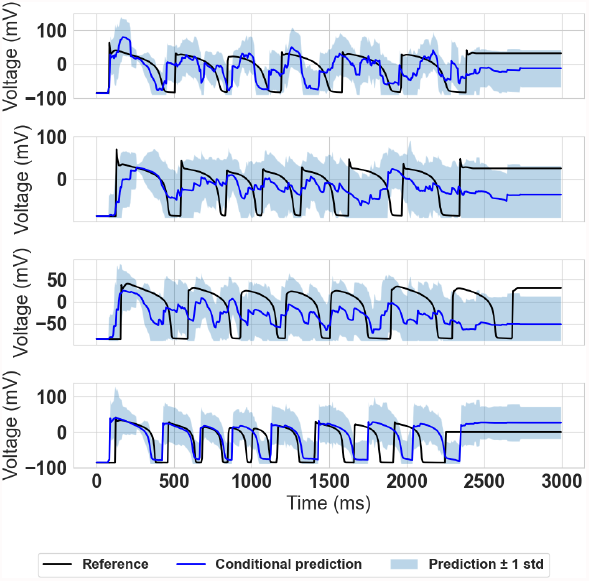
Conditional predictions for validation inputs in the VT regime.

**Fig. 12.**
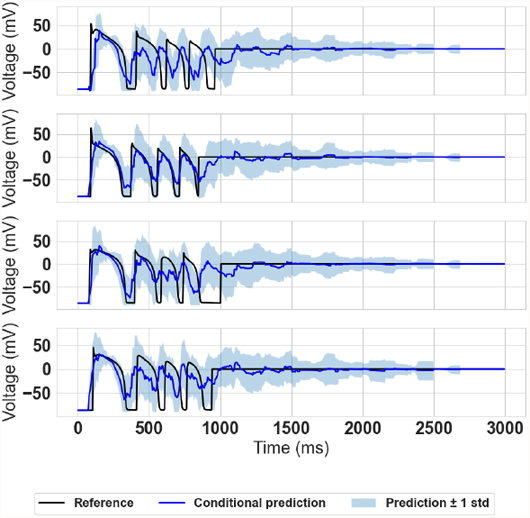
Conditional predictions for validation inputs in the non-VT regime.

## Methods

### B. Patient-specific 3D Model of Cardiac Electrophysiology

This study was approved by the IRB at Johns Hopkins University (IRB00190755). All procedures were conducted in accordance with relevant guidelines and regulations, and the retrospective analysis was performed on de-identified data. We utilized a patient’s 3D LGE-CMR data to reconstruct a fully resolved biventricular geometry capable of capturing the intricate right ventricular outflow tract (RVOT) and conal septum anatomy characteristic of repaired rToF hearts. The superior spatial resolution of 3D LGE-CMR allows for detailed segmentation of these regions, which are critical for modeling the re-entrant VT circuits commonly observed in rToF patients. The short-axis stack consisted of 71 slices (3.4 mm thickness, −1.7 mm inter-slice gap) with an in-plane resolution of 1.68 × 1.68 mm, obtained using a 1.5-T MRI scanner following the standardized clinical protocol. The images were resampled to an isotropic resolution of 0.35 × 0.35 × 0.35 mm to facilitate accurate segmentation. The right (RV) and left ventricles (LV) were delineated using a semi-automatic function of CardioViz3D, defining landmarks along the endocardial and epicardial borders extending to the pulmonary and tricuspid valves and the mitral annulus. This allowed for precise inclusion of the conal septum and RVOT.

Each myocardial voxel was classified as *normal, diffuse fibrosis*, or *dense scar* based on signal intensity using the *signal threshold to reference mean* method. Otsu thresholding was employed to segment the myocardium into high- and low-intensity regions, with separate thresholds computed for the RV and LV to account for differing intensity distributions. Voxels ≥5 SD above the mean were labeled as dense scar, and those between 3–5 SD as diffuse fibrosis. The ventricular septal defect patch, appearing as a low-intensity region surrounded by enhanced tissue, was manually corrected and labeled as scar. Cardiac fiber orientation vectors were assigned voxel-wise using a rule-based algorithm validated in prior studies.

The resulting geometry constitutes a detailed biventricular model faithfully representing a representative rToF patient. The geometric modeling pipline is summarized in Figure 13.

**Fig. 13.**
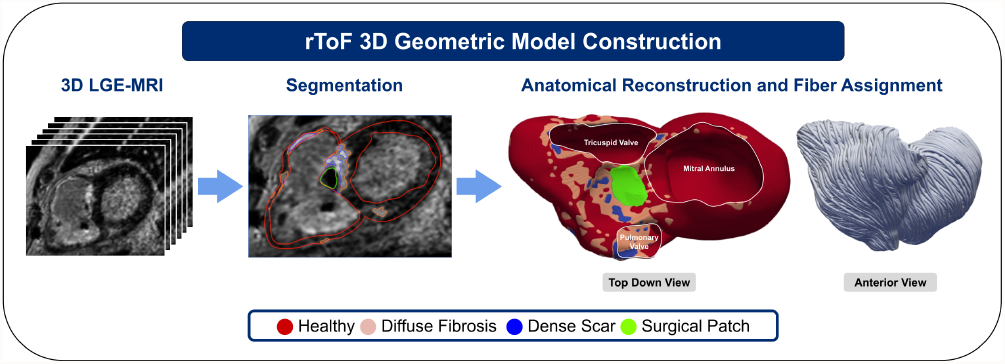
Schematic illustrating the geometric reconstruction of a rToF heart from LGE-CMR

#### B.1. Electrophysiological and tissue-level variability

Variability in cardiac anatomy, EP parameters, and tissue composition introduces significant uncertainty in predictive simulations of cardiac function. These uncertainties arise from multiple sources, including inter-individual differences in ionic conductances, heterogeneous fibrosis distribution, and modeling assumptions used during parameterization. Accurately representing such uncertainty is crucial for generating reliable DTs and for assessing their robustness under physiological and pathological perturbations. In this work, these uncertainties are explicitly incorporated into the generative manifold-based learning pipeline, enabling probabilistic sampling of diverse yet physiologically plausible heart models for downstream EP simulations.

##### Cell-Level Electrophysiological Variability

To account for inter-patient and intra-tissue variability in cell-level EP dynamics, we generated 100 variants of the ten Tusscher human ventricular myocyte model (34) by scaling the maximal conductances of six key ionic currents: fast sodium (INa), L-type calcium (ICaL), transient outward potassium (Ito), inward rectifier potassium (IK1), and the slow and fast delayed rectifier potassium currents (IKs, IKr). Scaling was applied only to the conductance parameters, preserving gating kinetics to reflect realistic biological variability in ion channel expression rather than structure. All currents except INa were scaled between 0.25 and 4 times baseline values, while INa was scaled between 0.25 and 3. Latin Hypercube Sampling (LHS) ensured uniform exploration of the parameter space. Each cell model was paced at 600 ms to steady-state, and action potential (AP) biomarkers (upstroke velocity, AP duration at 90% repolarization, and resting, plateau, and peak potentials) were compared to previously reported ranges (35) for validation.

Since fibrotic reomodeling has been shown to alter ion channel function (36), cell model variants for diffuse fibrosis were derived by applying remodeling factors according to the experimental data. Late sodium current (INaL), sodium-calcium exchanger (NCX), and ICaL were upregulated by factors of 1.07, 0.19, and 0.34, respectively, whereas IKr, IKs, Ito, IK1, and SERCA pump activity were downregulated by 0.34, 0.27, 0.85, 0.15, and 0.43, respectively. Gaussian noise (*µ* = 0, *σ* = 0.1) was added to scaling factors to reflect stochastic variability. The cell model variants were re-simulated to confirm that all the resulting APs remained physiologically plausible. Figure 14A compares the AP traces produced by the healthy and diffuse fibrosis cell model populations. On average, the fibrotic APs exhibit a longer AP duration, slower maximal upstroke velocity, and more resting membrane potential fluctuation.

**Fig. 14.**
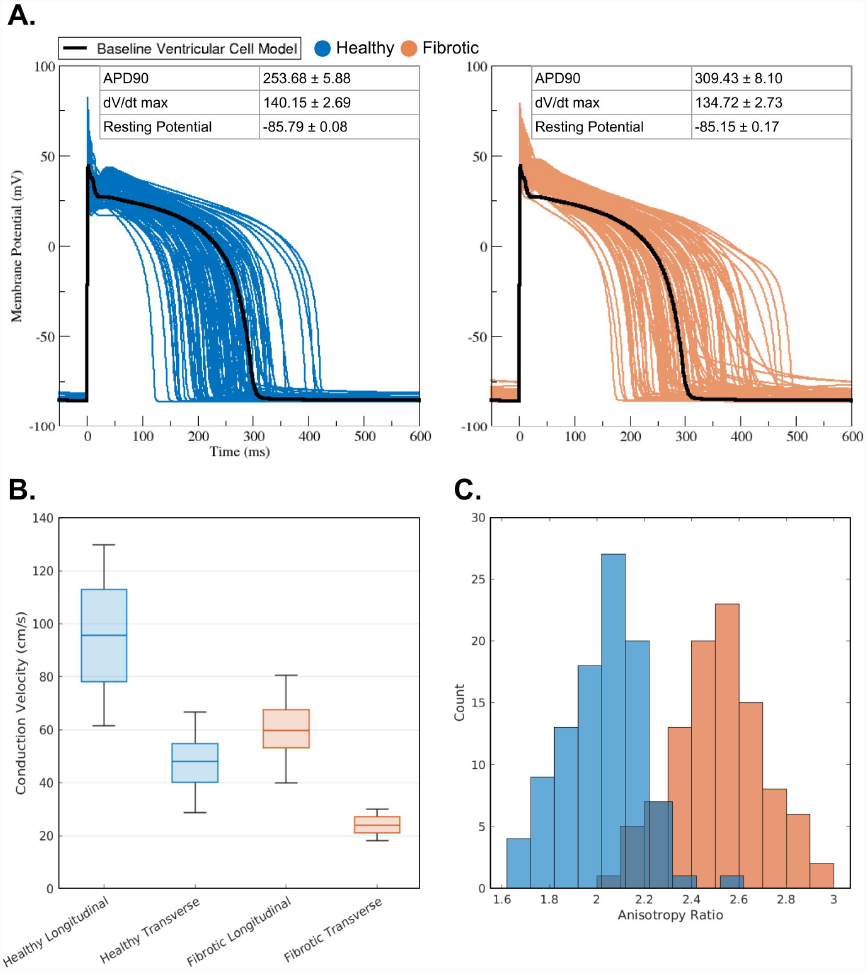
Healthy versus diffuse fibrosis Populations of Models: A.) Action potentials produced by varied ionic model parameters B.) Distributions of conduction velocities resulting from varied conductivity along and against fiber directions. C.) Frequency distribution of resulting anisotropy ratios (Longitudinal CV:Transverse CV)

Corresponding pairs of healthy and diffuse fibrosis model variants were mapped to their respective myocardial regions in the 3D geometric model, while dense scar voxels were modeled as electrical insulators.

##### Tissue-Level Variability

Cardiac conduction velocity (CV) also exhibits substantial inter-individual variability due to differences in tissue structure and myocardial fiber organization. The presence of fibrosis can further modify conduction properties, reducing overall CV and increasing CV anisotropy to varying degrees depending on the severity of tissue remodeling. To account for these uncertainties in our models, we generated 100 sets of conductivity tensors that represent electrical propagation along and across fiber directions for both healthy myocardium and diffuse fibrosis. The upper and lower extremes of CV values were first uniformly sampled from ranges corresponding to healthy longitudinal propagation and diffuse fibrosis transverse propagation.

- *CV*_*HL*_: 60–130 cm/s
- *CV*_*FT*_: 18–30 cm/s

Next, anisotropy ratios (AR) for both health regions and diffuse fibrosis were sampled from truncated normal distributions:

- *AR*_*Healthy*_ *∼ TN* (2.00, 0.023; 1.25, 3.00)
- *AR*_*GZ*_ *∼ TN* (2.50, 0.04; 1.25, 3.75)

where *TN* (*µ, σ*^2^; *a, b*) denotes a normal distribution with mean *µ*, variance *σ*^2^, truncated to the interval [*a,b*]. The sampled ratios were then used to back-calculate the healthy transverse (*CV*_*HT*_) and diffuse fibrosis longitudinal (*CV*_*FL*_) conduction velocities.

- 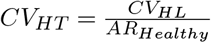
- *CV*_*FL*_ = *CV*_*FT*_ × *AR*_*GZ*_

Figure 14B illustrates the resulting CV distributions derived from the sampling procedure. The box plots highlight the expected hierarchy of conduction speeds, where conduction was fastest along the healthy longitudinal direction, slowest along the diffuse fibrosis transverse direction, and intermediate for the remaining two cases. These distributions are in line with previous modeling studies of rToF (37). The accompanying histograms in Figure 14C compare the AR distributions for healthy tissue and diffuse fibrosis, demonstrating a rightward shift in the diffuse fibrosis AR distribution that reflects increased directional disparity in conduction due to remodeling effects.

Finally, conductivities corresponding to the desired CVs were determined by simulating uniaxial propagation along a 20 mm × 7 mm × 3 mm slab at 400 *µ*m resolution. Fiber orientation was aligned with the *x*-axis for longitudinal and with the *y*-axis for transverse tuning. Calibrated conductivity tensors were then assigned voxel-wise to the tissue regions in the 3D geometric model.

#### B.2. Ventricular Tachycardia Induction Simulations

The modeling pipeline yielded a cohort of 100 rToF DT variants capturing the uncertainty arising from unknown EP properties within a common cardiac geometry. For each DT, a rapid pacing protocol (4) was used to test for VT inducibility. Six S1 stimuli were delivered to the conal septum at a basic cycle length of 600 ms, followed by up to three premature (S2–S4) stimuli with progressively shorter coupling intervals. A simulation was labeled as *inducible* if, following cessation of pacing, the model exhibited 2 s of periodic depolarization and repolarization due to self-sustaining wavefront propagation. Figure 1 illustrates two example VTs arising from different model parameter set.

All simulations were performed using internally developed software for solving cardiac electrophysiology models. Using five Intel Xeon w7-3445 cores with 24 GB of allocated memory and a Nvidia RTX A4500 GPU, the wall time for running 100 simulations was approximately 100 hours.

### C. Double Diffusion Maps Probabilistic Learning on Manifolds

Uncertainty in cardiac DTs arises from the non-linear coupling of EP, anatomical, and biophysical model parameters, which often reside on a low-dimensional manifold embedded in a high-dimensional parameter space. Traditional UQ methods treat these parameters as independent random variables, neglecting the geometric dependencies that constrain physiologically plausible cardiac configurations. To overcome this limitation, we employ DD-PLoM (32), a generative manifold learning framework introduced by the authors, to learn the joint probability density over cardiac observables and model parameters, enabling the characterization of statistical dependencies across a large number of variables in patient-specific cardiac. Let **Q** denote the quantity of interest (e.g., transmambrane potential signal) and **W** the set of observed variables (cell-level electrophysiological and tissue properties associated with ventricular tachycardia). In our setting, each data point consists of the concatenated vector **X** = [**W, R**] ∈ ℝ^*N*×*d*^. Our goal is to learn the joint distribution *p*(**W, R**) and perform conditional sampling. Next, we discuss the basic ingredients of DD-PLoM and non-parametric conditioning for sampling and UQ.

#### C.1. Diffusion Maps manifold for dimensionality reduction

Let **X**^⊺^ ≡ [*x*_*d*_] = [*x*^(*1*)^, …, *x*^*(N)*^] ∈ ℝ^*d×N*^ be the data matrix, where each column *x*^(*j*)^ ∈ ℝ^*n*^. After standard preprocessing (e.g., centering, scaling), we construct a diffusion kernel

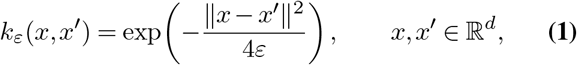

with scale parameter *ε >* 0, and define the kernel matrix

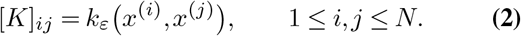

A good practical initial guess for the diffusion maps kernel scale parameter is the squared median pairwise distance between data points. Diffusion Maps normalization introduces matrices:

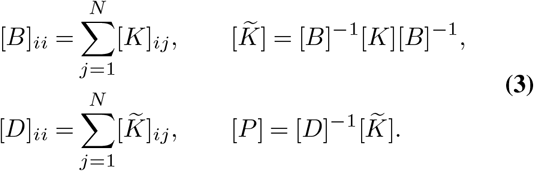

as well as the symmetric matrix:

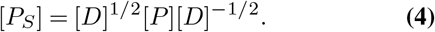

The solution of the eigenvalue problem

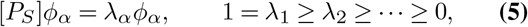

provides a set of scaled diffusion coordinates

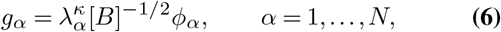

for a chosen diffusion time *κ* ∈ ℕ. Selecting the first *M* co-ordinates gives

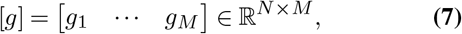

However, not all coordinates *g*_*α*_ are equally informative. DD-PLoM distinguishes non–harmonic (intrinsic) coordinates from harmonic ones using a localized regression criterion: for each eigenvector we attempt to express it as a nonlinear function of a small subset of leading coordinates; eigenvectors with large residuals are labeled non–harmonic. To retain only the *m* non–harmonic coordinates that correspond to a new, unique eigendirection, the authors in (38) define a normalized leave-one-out cross-validation error based on local linear regression:

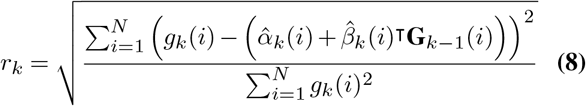

Here, **G**_*k*−1_(*i*) = [*g*_1_(*i*),…, *g*_*k*−1_(*i*)]^⊺^ is the vector of the previous (*k*− 1) eigenvectors at point *i*, and the coefficients 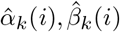 are obtained from locally weighted linear regression. A small *r*_*k*_ suggests that *g*_*k*_ is a harmonic (i.e., repeated eigen-direction), while a large *r*_*k*_ indicates a unique new direction.

The leading non-harmonic eigenvectors of the Markov operator define a reduced set of diffusion coordinates [*g*_*m*_] = [*g*^(1)^, …, *g*^*(m)*^] ∈ ℝ^*N*×*m*^, which parametrize the intrinsic geometry of the data manifold. We need to highlight here that, the choice of kernel function *k*(*·,·*) determines the geometry encoded in the affinity matrix and directly influences the latent coordinates. In this work, we adopt the Gaussian kernel, owing to its smoothness, locality, and asymptotic link to the Laplace–Beltrami operator (27).

#### C.2. Geometric Harmonics

A key challenge when working with points on manifolds is lifting them back to the ambient space: given a new latent point *g*_new_ ∈ ℝ^*m*^, we must reconstruct *x*_new_ ∈ ℝ^*n*^ consistent with the manifold geometry and the data. To do this we can use Geometric Harmonics (GH), a kernel-based approach for extending functions defined on a dataset to new, unseen point, introduced by Coifman and Lafon (39). Let 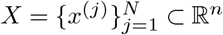 and 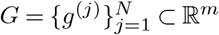. Consider a scalar function *f*: *G* → ℝ defined on the latent sample set, with values *f* ^(*j*)^ = *f* (*g*^(*j*)^). We approximate *f* by projection onto the latent harmonics:

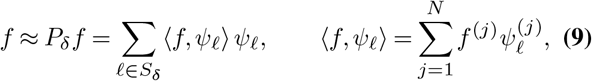

where *S*_*δ*_ = {*ℓ*: *σ*_*ℓ*_ ≥*δσ*_0_} is a spectral truncation set and *δ* ∈ (0, 1) controls the smoothness of the approximation (large *δ* retains only the smoothest modes). To evaluate *f* at a new latent point *g*_new_, we first extend each eigenvector *ψ*_*ℓ*_ via a Nyström–type formula:

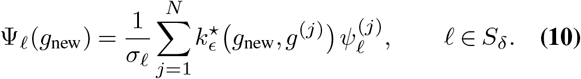

The GH extension of *f* at *g*_new_ is then

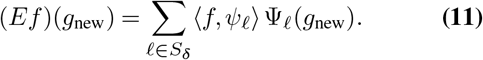

We apply this construction componentwise to the vector–valued map

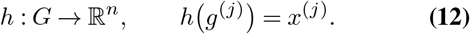

Each physical component (e.g., potential at a mesh node, or a particular ECG channel at a given time) is treated as a scalar function on *G* and extended via Eq. (11). The resulting operator

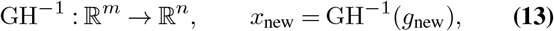

acts as a manifold–aware decoder: it lifts a latent cardiac state *g*_new_ to a high–dimensional cardiac realization *x*_new_ consistent with the learned manifold.

#### C.3. Function Representation on the Manifold

To accurately represent functions on the learned manifold, it becomes necessary to perform a second Diffusion Maps step directly in the intrinsic space (40). This additional step is necessary because, in the initial embedding, higher-order eigenvectors are truncated to obtain a parsimonious representation. The discarded modes still carry essential information for accurately lifting functions back to the ambient space. By recomputing Diffusion Maps in the intrinsic coordinates, we recover a functional basis that enables consistent and smooth lifting.

Given 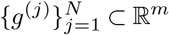, we define a latent kernel

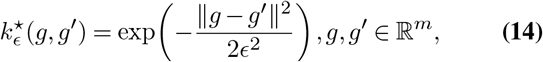

and the associated matrix 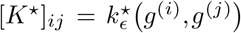. We compute the first *L* eigenpairs [*K*^⋆^]*ψ*_*ℓ*_ = *σ*_*ℓ*_*ψ*_*ℓ*_, with *ℓ* = 0,…, *L* − 1, and collect the leading eigenvectors into

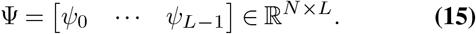

These {*ψ*_*ℓ*_} form a data–adapted basis of latent harmonics on the finite point cloud {*g*^*(j)*^}. Intuitively, they play a role analogous to Laplace–Beltrami eigenfunctions on the manifold, providing multi–scale building blocks for smooth functions defined on this manifold.

#### C.4. Latent density and full–order ISDE

In the intrinsic space ℝ^*m*^, we estimate the probability density of the latent variables using a kernel density estimate:

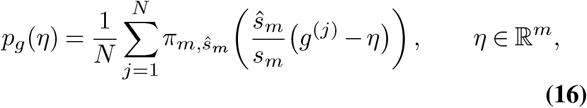

where *π*_*m,ŝm*_ is an *m*–dimensional Gaussian kernel and *s*_*m*_, *ŝ*_*m*_ are bandwidth parameters chosen so that the transformed latent variables have zero mean and identity covariance. This density captures the intrinsic variability of the cardiac states on the learned manifold. Following (32), we then construct an Itô stochastic dynamical system in latent space whose invariant measure is *p*_g_ Let

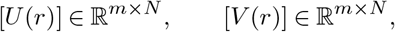

denote the latent position and velocity processes, where *r* ≥0 is a fictitious “time” parameter for sampling (not the physical time of the cardiac dynamics). We consider the ISDE

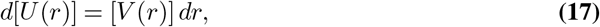

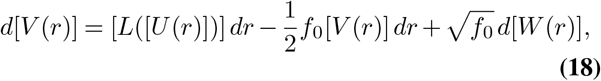

where:

- *f*_0_ *>* 0 is a damping parameter,
- [*W* (*r*)] is a matrix–valued Wiener process in ℝ^*m*×*N*^,
- [*L*([*U*])] is defined via the gradient of the log–density,
- 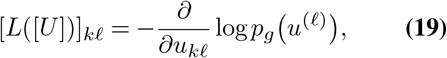
- with *u*^(*ℓ*)^ the *ℓ*–th column of [*U*].

By construction, the invariant measure of Eq. (17)–Eq. (18) coincides with the product distribution induced by *p*_g_. After an initial relaxation phase, samples of [*U* (*r*)] therefore explore the latent manifold according to the learned distribution.

### D. Non-parametric conditional sampling

In many scientific and engineering settings, predictions of the quantity of interest must be made conditional on partially observed inputs, rather than in an unconditional sense. In particular, we are often interested not only in the average response, but in the full distribution of outcomes that are consistent with a given set of observed variables. This necessitates accurate approximations of conditional distributions, especially in high-dimensional and data-sparse regimes where classical parametric approaches are either intractable or overly restrictive.

Given observations **W** = **w**_0_, the objective is to infer **Q** through the conditional distribution *p*(**Q** | **W** = **w**_0_). To enable this, we learn a joint probability distribution *p*(**Q, W**) over all variables, and inference is carried out through conditioning on the learned joint distribution using a non-parametric approach. Let 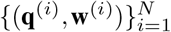 denote samples drawn from the learned joint distribution, where **q**^(*i*)^ and **w**^(*i*)^ correspond to realizations of **Q** and **W**, respectively.

The conditional expectation of **Q** given **W** = **w**_0_ is approximated using a kernel-based estimator:

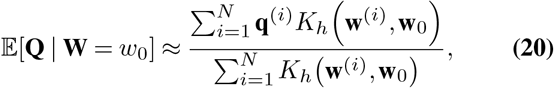

where *K*_*h*_(*·,·*) is a positive kernel function with bandwidth parameter *h*. In this work, we employ a Gaussian kernel of the form

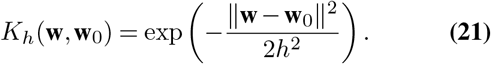

This estimator assigns higher weights to samples whose conditioning variables **w**^(*i*)^ are closer to the target value **w**_0_, thereby enabling local approximation of the conditional distribution without assuming a parametric form. Beyond conditional expectations, the full conditional distribution can be approximated using weighted empirical measures. For a scalar quantity of interest *Q*_*j*_, the conditional cumulative distribution function (CDF) is estimated as

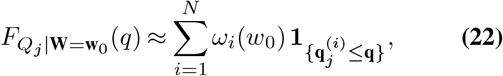

where the normalized weights are given by

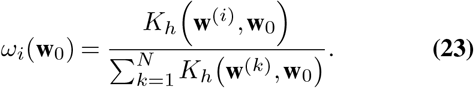

The corresponding conditional probability density function (PDF) can be obtained by differentiation or via kernel density estimation using the same weights. Furthermore, conditional samples can be generated by resampling from the set {**q**^(*i*)^} with probabilities {*ω*_*i*_(**w**_0_)}, yielding realizations that approximate samples from *p*(**Q** | **W** = **w**_0_). This enables the construction of conditional ensembles for downstream analysis, including uncertainty characterization and classification tasks. This non-parametric conditioning approach is particularly well-suited to the present setting, where the learned joint distribution is defined implicitly through samples on a nonlinear manifold and does not admit a closed-form representation. By operating directly on the generated samples, the method avoids restrictive parametric assumptions while, by construction, preserving the intrinsic geometric structure of the data.

#### Local Gaussian approximation

In addition to the non-parametric empirical approximation, we also consider a local Gaussian representation of the conditional distribution. Specifically, the conditional distribution *p*(**Q** | **W** = **w**_0_) is approximated by a Gaussian distribution

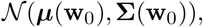

here the mean and covariance are estimated using the same kernel weights {*ω*_*i*_(**w**_0_)} as

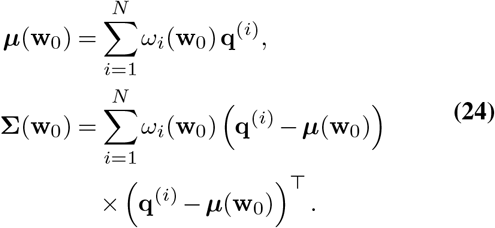

This approximation provides a convenient parametric surrogate for the conditional distribution, enabling comparison between empirical sampling and Gaussian assumptions, as illustrated in the figures.

#### Random conditional sampling

While the weighted empirical construction above provides a deterministic approximation of conditional statistics, we need the ability to draw random realizations from the conditional distribution *p*(**Q** | **W** = **w**_0_). To this end, we interpret the normalized weights 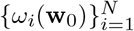 as a discrete probability measure supported on the samples 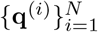, and define a random conditional draw **Q**^*∗*^ via

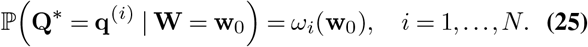

Equivalently, **Q**^*∗*^ can be generated by sampling an index

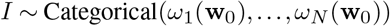

and setting **Q**^*∗*^ = **q**^(*i*)^. Repeating this procedure yields an ensemble 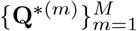 of conditionally consistent realizations. The need for such random conditional sampling is twofold. First, it enables the propagation of uncertainty under conditioning, allowing us to characterize not only mean responses but also variability, tail behavior, and rare events associated with a given **w**_0_. This is particularly critical in high-dimensional, data-sparse regimes, where deterministic summaries (e.g., conditional expectations) may obscure multimodality or heteroscedastic effects. Second, random conditional samples provide a practical mechanism for tasks such as classification, risk quantification, and decision-making, where Monte Carlo estimates of functionals of **Q** are required. Importantly, this sampling strategy remains fully non-parametric and is consistent with the learned joint distribution, as it relies solely on the empirical support of generated samples and their locality in **W**-space. As a result, it preserves the intrinsic geometry and correlations encoded in the data, avoiding artifacts that could arise from imposing an explicit parametric form on *p*(**Q** | **W**). In practice, this yields conditionally coherent ensembles that reflect both the learned manifold structure and the uncertainty induced by limited observations.

## Discussion

In this study we combined a high–fidelity, patient–specific model of rToF with a geometry–aware generative learning framework (DD–PLoM) to characterize how uncertainty in EP and tissue properties affects ventricular arrhythmia inducibility. Throughout, we use the terms VT / non–VT and arrhythmic / non–arrhythmic interchangeably to refer to models that did or did not sustain reentry under the programmed stimulation protocol. Below we discuss the main mechanistic and methodological implications of our findings, their potential clinical relevance, and limitations that motivate future work.

### E. Mechanistic insights from uncertain model parameters

We first examined how uncertainty in model parameters influenced whether simulations developed VT. Although all models used the same heart anatomy and scar distribution, varying cellular and tissue properties produced a wide range of outcomes, from non-arrhythmic behavior to sustained reentry.

At the cellular level, simulations that developed VT tended to occupy different regions of the sampled parameter space than those that did not. In particular, models with VT more often combined increased excitability with changes in recovery properties, reflecting shifts in how electrical signals are initiated and maintained. These trends should not be interpreted as identifying a single causal parameter. Rather, they indicate directions in parameter space that make the system more sensitive to reentry within the ranges explored here. Different modeling choices or parameter ranges could change the importance of individual variables while preserving the overall dependence on combined effects.

At the tissue level, VT was more closely associated with slowed electrical propagation in surrounding healthy regions than with large differences inside fibrotic tissue itself. This highlights an important distinction between structure and function. While scar and surgical patches define possible pathways for reentry, whether these pathways actually sustain VT depends on how electrical activity travels through the surrounding tissue.

Taken together, these findings show that arrhythmia risk cannot be understood from anatomy alone or from a single “best-fit” set of parameters. Instead, VT emerges from the interaction of multiple uncertain properties acting together. This motivates uncertainty-aware modeling approaches that describe a range of possible behaviors, rather than relying on a deterministic prediction from a single parameter set, when using digital heart models to assess arrhythmia risk.

### F. Performance of conditional sampling and uncertainty quantification

A central objective of the proposed framework is to *enable accurate conditional inference of cardiac responses given partially observed or prescribed electrophysiological parameters*. Building upon the learned joint distribution *p*(**W, R**), we perform non-parametric conditional sampling to approximate *p*(**R** | **W** = **w**_0_) and quantify the associated uncertainty. The results demonstrate that the conditional distributions capture both the mean behavior and variability of the transmembrane potential trajectories. The conditional mean responses closely track the reference simulations across all representative parameter configurations, accurately reproducing key electrophysiological features such as depolarization upstroke, oscillatory dynamics, and repolarization phases. Importantly, the associated uncertainty bands are not uniform in time: variability increases during highly nonlinear phases of the dynamics (e.g., reentry onset and inter-spike intervals), while remaining tighter during more stable phases such as depolarization and plateau. This behavior reflects the intrinsic sensitivity of the cardiac dynamics to parameter perturbations.

The quality of the learned conditional distribution is further assessed through random conditional sampling. The generated samples preserve the qualitative and quantitative structure of the reference trajectories, including activation timing, amplitude, and oscillatory patterns. The agreement improves for conditioning points that lie in regions of higher data density, where the kernel-based weighting emphasizes nearby samples in parameter space. As the distance from the conditioning point increases, discrepancies in amplitude and timing may emerge, reflecting increased epistemic uncertainty due to sparse data coverage. Nevertheless, even in these regimes, the generated samples remain physiologically plausible, indicating that the learned manifold captures the admissible space of cardiac dynamics.

To quantitatively evaluate predictive performance, we consider three complementary metrics: coverage, tail probability, and root mean squared error (RMSE). Coverage measures the fraction of time instances for which the reference trajectory lies within the predicted conditional uncertainty band, providing a time-resolved assessment of reliability. Tail probability evaluates the global plausibility of the reference trajectory under the learned conditional distribution by comparing its deviation from the conditional mean to the distribution of deviations of sampled trajectories; low values indicate that the reference lies in a low-density region. RMSE quantifies the discrepancy between the conditional mean prediction and the reference trajectory, capturing the bias of the estimator. Across validation cases, the conditional predictions exhibit consistently high coverage (typically close to or above 0.9), moderate tail probabilities, and low RMSE values. These results indicate that the learned conditional distribution not only provides accurate central predictions but also produces uncertainty estimates that are well-calibrated and statistically consistent with the observed data. In particular, high coverage combined with moderate tail probabilities suggests that the reference trajectories lie within high-probability regions of the learned distribution, while the relatively low RMSE confirms that the conditional mean captures the dominant dynamics. However, the validation analysis also reveals limitations inherent to the data-sparse, high-dimensional setting. In certain cases, reduced coverage and lower tail probability are observed despite low RMSE, indicating that the uncertainty bands may underestimate variability in specific regions of the input space. This behavior is consistent with the limited number of training samples (39 VT and 61 non-VT cases) in a 596-dimensional space, where local neighborhoods are poorly populated and conditional estimation becomes more challenging.

Overall, these results demonstrate that the proposed geometry-aware generative framework enables accurate, uncertainty-aware conditional inference of cardiac dynamics. By combining manifold-based learning with non-parametric conditioning, the approach captures both the central tendencies and variability of the system, providing a robust alternative to classical UQ methods that would otherwise require a prohibitively large number of forward simulations.

### G. Clinical implications and potential applications

From potential clinical applications standpoint, the proposed framework offers several potential advantages. First, it provides a principled way to incorporate uncertainty into digital–twin simulations without requiring thousands of expensive, full–order model runs. In this study, approximately one hundred high–fidelity simulations were sufficient to train DD–PLoM; thereafter, tens of thousands of additional virtual realizations could be generated at negligible computational cost. This may enable practical exploration of “what–if” scenarios and sensitivity analyses for individual patients, for example quantifying how plausible variability in conductivities, fibrosis properties, or ionic remodelling might alter VT inducibility or ablation outcomes.

Second, by learning the statistical relationship between parameter combinations and resulting EP behavior, the framework can be integrated into inverse modeling or data-assimilation workflows. In this setting, the generative model could be used to constrain patient-specific model fitting to parameter combinations that are consistent with known physiology and with the simulated arrhythmic and non-arrhythmic behaviors. For example, when fitting DT models to clinical data such as CV slowing from intra-procedural electroanatomic maps, inference could be restricted to regions of parameter space that lie on the learned manifold, rather than exploring an unrestricted high-dimensional space. This constraint has the potential to improve both computational efficiency and physiological plausibility of patient-specific model calibration.

Third, separating the learning of the parameter–response structure from the subsequent sampling stage enables reuse of the trained generative model as new clinical questions arise. Once the relationship between parameter combinations and EP behavior has been learned for a given anatomical scenario, the same model can be leveraged to explore alternative stimulation protocols, pharmacological perturbations, or pacing strategies without retraining. In practice, this allows efficient resampling of the learned parameter space and selective re-simulation of configurations that are predicted to be arrhythmogenic. More broadly, this approach could support the construction of libraries of learned parameter manifolds associated with distinct scar patterns or ventricle shapes, providing a foundation for patient-stratified DT analyses.

### H. Limitations

Several limitations must be acknowledged when interpreting these results. First, the present study focuses on a single representative anatomy of rToF. Although the uncertainty in ionic conductances and conduction properties was substantial, the anatomical variability seen across patients with rToF is not yet represented. Extending the framework to multiple anatomies and scar patterns will be important to ensure that the learned manifolds generalize beyond a single structural template. Second, we considered a specific set of ionic currents and conduction parameters, informed by prior work, but other sources of variability (e.g. autonomic tone, loading conditions, regional differences in coupling or gap junction expression) were not explicitly modeled. These factors may further modulate VT risk and could be incorporated in future versions of the DTs. Third, although DD–PLoM reproduces the distributions of simulated parameters and voltage responses, we have not yet validated its predictions directly against clinical data (such as ECG morphologies or invasive mapping) in a larger patient population. Our present analysis should therefore be viewed as a proof of concept in a controlled *in silico* setting. Translational use will require careful calibration and validation against prospective clinical measurements. Fourth, the performance of diffusion–maps–based methods depends on several hyperparameters (kernel bandwidths, number of retained eigenmodes, density–estimation bandwidths). In this work, these were chosen based on heuristics and exploratory analyses. Systematic strategies for hyperparameter selection and uncertainty quantification within the manifold–learning step itself will be important for robust clinical deployment. Finally, DD–PLoM is an *unsupervised generative model*: it captures the joint variability of parameters and responses but does not, by itself, provide a predictive classifier for VT versus non–VT outcomes. In practice, our framework would likely be combined with additional supervised models or decision rules (for example, logistic regression or machine–learning classifiers trained on latent variables) to estimate individualized VT risk or to guide ablation targeting.

### I. Future directions

Future work will focus on three main directions. First, we plan to extend the analysis to multi–patient datasets and to other structural substrates such as non–ischaemic cardiomyopathies, with the goal of identifying how the learned manifolds and arrhythmia–associated regions change with anatomy and disease. Second, we aim to couple DD–PLoM to clinical observables, using the generative model as a prior in Bayesian inversion of ECGs or intracardiac electrograms to patient–specific parameter distributions. Third, we envision exploiting the efficiency of the generative surrogate to explore alternative stimulation protocols, anti–arrhythmic drug effects, or device settings within the same framework. In summary, the present study demonstrates that geometry–aware generative learning can faithfully reproduce the complex, coupled uncertainty in a high–fidelity cardiac DT and can do so with far fewer full electrophysiology simulations than traditional Monte Carlo approaches. For clinicians, the key message is that such methods offer a way to move from a single deterministic DT prediction to a probabilistic description of arrhythmic risk that explicitly accounts for uncertainty in tissue properties and EP modeling.

## Conclusions

In this work we combined a detailed, patient–specific model of rToF with a geometry–aware generative learning framework to study how uncertainty in EP and tissue properties shapes the propensity for VT. Throughout, we used the terms VT / non–VT and arrhythmic / non–arrhythmic interchangeably to distinguish simulations that did or did not sustain reentry under a clinically motivated stimulation protocol Our analysis showed that, even for a fixed anatomy, realistic variability in ionic conductances and conduction properties can move the system between clearly non–arrhythmic behaviour and stable, sustained VT. Arrhythmic simulations were enriched in parameter combinations that prolong repolarization and increase repolarization heterogeneity, together with highly anisotropic, slowed conduction in diffuse fibrosis surrounding scar and surgical patch regions. These mechanisms are consistent with clinical observations of VT circuits in repaired Tetralogy of Fallot and highlight the importance of representing parameter uncertainty when interpreting digital–twin predictions.

Methodologically, we demonstrated that DD–PLoM can learn a compact, low–dimensional manifold that captures the joint variability of parameters and electrophysiological responses, rather than fitting each parameter distribution separately. Once trained, the generative model was able to produce thousands of additional virtual realizations that closely matched the reference distributions of parameters, action–potential waveforms, and time–resolved voltage densities in both arrhythmic and non–arrhythmic groups. Beyond reproducing marginal statistics, the framework enables non–parametric conditional sampling, allowing us to generate response trajectories conditioned on specific parameter configurations. The resulting conditional predictions accurately recover key dynamical features of the reference simulations, while the associated uncertainty bands reflect the time–dependent variability of the underlying electrophysiological processes. Quantitative evaluation using coverage, tail probability, and RMSE demonstrates that the learned conditional distributions are both well–calibrated and predictive, capturing the dominant temporal structure of the transmembrane potential while providing meaningful measures of uncertainty. From a practical standpoint, the proposed framework provides a route toward DTs that report probabilistic, rather than purely deterministic, assessments of arrhythmic risk, supporting “what–if” analyses, sensitivity studies, and future inverse–problem formulations that assimilate clinical data under explicit uncertainty. Future work will extend this approach to multiple anatomies and disease substrates, integrate clinical measurements (e.g. ECGs and electroanatomic maps) into the generative framework, and explore the use of DD–PLoM–based surrogates to guide patient–specific VT risk stratification and ablation planning. Overall, our results suggest that geometry–aware generative learning is a promising tool for bringing uncertainty–aware, clinically meaningful cardiac DTs closer to routine use.

## ACKNOWLEDGEMENTS

This research was supported by the National Science Foundation (under Grant No 2436738) and the National Heart, Lung, and Blood Institute, National Institutes of Health (R01HL166759, R01HL174440, T32HL007024). The funders played no role in study design, data collection, analysis and interpretation of data, or the writing of this manuscript. This material is also based upon work supported by the National Science Foundation Graduate Research Fellowship Program (DGE2139757). Any opinions, findings, and conclusions or recommendations expressed in this material are those of the author(s) and do not necessarily reflect the views of the National Science Foundation.

## Bibliography

1. Hermenegild J. Arevalo, Fijoy Vadakkumpadan, Eliseo Guallar, Alexander Jebb, Peter Malamas, Katherine C. Wu, and Natalia A. Trayanova. Arrhythmia risk stratification of patients after myocardial infarction using personalized heart models. 7(1):11437. ISSN 2041-1723. doi: 10.1038/ncomms11437.

2. Yingnan Zhang, Kelly Zhang, Adityo Prakosa, Cynthia James, Stefan L Zimmerman, Richard Carrick, Eric Sung, Alessio Gasperetti, Crystal Tichnell, Brittney Murray, Hugh Calkins, and Natalia A Trayanova. Predicting ventricular tachycardia circuits in patients with arrhythmogenic right ventricular cardiomyopathy using genotype-specific heart digital twins. 12:RP88865. ISSN 2050-084X. doi: 10.7554/eLife.88865.

3. Eric Sung, Adityo Prakosa, Shijie Zhou, Ronald D. Berger, Jonathan Chrispin, Saman Nazarian, and Natalia A. Trayanova. Fat infiltration in the infarcted heart as a paradigm for ventricular arrhythmias. 1(10):933–945. ISSN 2731-0590. doi: 10.1038/s44161-022-00133-6.

4. Adityo Prakosa, Hermenegild J. Arevalo, Dongdong Deng, Patrick M. Boyle, Plamen P. Nikolov, Hiroshi Ashikaga, Joshua J. E. Blauer, Elyar Ghafoori, Carolyn J. Park, Robert C. Blake, Frederick T. Han, Rob S. MacLeod, Henry R. Halperin, David J. Callans, Ravi Ranjan, Jonathan Chrispin, Saman Nazarian, and Natalia A. Trayanova. Personalized virtual-heart technology for guiding the ablation of infarct-related ventricular tachycardia. 2(10):732–740. ISSN 2157-846X. doi: 10.1038/s41551-018-0282-2.

5. Haibo Ni, Stefano Morotti, and Eleonora Grandi. A heart for diversity: Simulating variability in cardiac arrhythmia research. 9. ISSN 1664-042X. doi: 10.3389/fphys.2018.00958.

6. Zhiyong Hu, Dongping Du, and Yuncheng Du. Generalized polynomial chaos-based uncertainty quantification and propagation in multi-scale modeling of cardiac electrophysiology. 102:57–74. ISSN 0010-4825. doi: 10.1016/j.compbiomed.2018.09.006.

7. Pras Pathmanathan, Suran K. Galappaththige, Jonathan M. Cordeiro, Abouzar Kaboudian, Flavio H. Fenton, and Richard A. Gray. Data-driven uncertainty quantification for cardiac electrophysiological models: Impact of physiological variability on action potential and spiral wave dynamics. 11. ISSN 1664-042X. doi: 10.3389/fphys.2020.585400.

8. Eric Schulz, Maarten Speekenbrink, and Andreas Krause. A tutorial on gaussian process regression: Modelling, exploring, and exploiting functions. Journal of mathematical psychology, 85:1–16, 2018.

9. Stefania Fresca, Andrea Manzoni, Luca Dedè, and Alfio Quarteroni. Deep learning-based reduced order models in cardiac electrophysiology. PloS one, 15(10):e0239416, 2020.

10. Lucian Itu, Puneet Sharma, Vwrel Mihalef, Ali Kamen, Constantin Suciu, and Dorm Lomaniciu. A patient-specific reduced-order model for coronary circulation. In 2012 9th IEEE international symposium on biomedical imaging (ISBI), pages 832–835. IEEE, 2012.

11. Elena Sabdy Martinez, Beatrice Moscoloni, Matteo Salvador, Fanwei Kong, Mathias Peirlinck, and Alison Lesley Marsden. Full-field surrogate modeling of cardiac electrophysiology encoding geometric variability. 448:118444. ISSN 0045-7825. doi: 10.1016/j.cma.2025.118444.

12. Diederik P Kingma. Auto-encoding variational bayes. arXiv preprint 1312.6114, 2013.

13. Danilo Jimenez Rezende, Shakir Mohamed, and Daan Wierstra. Stochastic backpropagation and approximate inference in deep generative models. In International conference on machine learning, pages 1278–1286. PMLR, 2014.

14. George Papamakarios, Eric Nalisnick, Danilo Jimenez Rezende, Shakir Mohamed, and Balaji Lakshminarayanan. Normalizing flows for probabilistic modeling and inference. Journal of Machine Learning Research, 22(57):1–64, 2021.

15. Ian Goodfellow, Jean Pouget-Abadie, Mehdi Mirza, Bing Xu, David Warde-Farley, Sherjil Ozair, Aaron Courville, and Yoshua Bengio. Generative adversarial networks. Communications of the ACM, 63(11):139–144, 2020.

16. Mehdi Mirza. Conditional generative adversarial nets. arXiv preprint 1411.1784, 2014.

17. Ling Yang, Zhilong Zhang, Yang Song, Shenda Hong, Runsheng Xu, Yue Zhao, Wentao Zhang, Bin Cui, and Ming-Hsuan Yang. Diffusion models: A comprehensive survey of methods and applications. ACM computing surveys, 56(4):1–39, 2023.

18. Yang Song and Stefano Ermon. Generative modeling by estimating gradients of the data distribution. In H. Wallach, H. Larochelle, A. Beygelzimer, F. d’ Alché-Buc, E. Fox, and R. Garnett, editors, Advances in Neural Information Processing Systems, volume 32. Curran Associates, Inc.

19. Yang Song, Jascha Sohl-Dickstein, Diederik P. Kingma, Abhishek Kumar, Stefano Ermon, and Ben Poole. Score-based generative modeling through stochastic differential equations, 2021.

20. Yang Song and Stefano Ermon. Improved techniques for training score-based generative models. Advances in neural information processing systems, 33:12438–12448, 2020.

21. Zhifeng Kong, Wei Ping, Jiaji Huang, Kexin Zhao, and Bryan Catanzaro. Diffwave: A versatile diffusion model for audio synthesis. arXiv preprint 2009.09761, 2020.

22. Chenhao Niu, Yang Song, Jiaming Song, Shengjia Zhao, Aditya Grover, and Stefano Ermon. Permutation invariant graph generation via score-based generative modeling. In International Conference on Artificial Intelligence and Statistics, pages 4474–4484. PMLR, 2020.

23. Joseph L Watson, David Juergens, Nathaniel R Bennett, Brian L Trippe, Jason Yim, Helen E Eisenach, Woody Ahern, Andrew J Borst, Robert J Ragotte, Lukas F Milles, et al. Broadly applicable and accurate protein design by integrating structure prediction networks and diffusion generative models. BioRxiv, pages 2022–12, 2022.

24. Charles Fefferman, Sanjoy Mitter, and Hariharan Narayanan. Testing the manifold hypothesis. Journal of the American Mathematical Society, 29(4):983–1049, 2016.

25. Mukund Balasubramanian and Eric L Schwartz. The isomap algorithm and topological stability. Science, 295(5552):7–7, 2002.

26. Sam T Roweis and Lawrence K Saul. Nonlinear dimensionality reduction by locally linear embedding. science, 290(5500):2323–2326, 2000.

27. RR Coifman and S Lafon. Diffusion maps. Applied and Computational Harmonic Analysis, 21(1):5–30, 2006.

28. Christian Soize and Roger Ghanem. Data-driven probability concentration and sampling on manifold. Journal of Computational Physics, 321:242–258, 2016.

29. Christian Soize, Roger Ghanem, Cosmin Safta, Xun Huan, Zachary P Vane, J Oefelein, Guilhem Lacaze, Habib N Najm, Qi Tang, and X Chen. Entropy-based closure for probabilistic learning on manifolds. Journal of Computational Physics, 388:518–533, 2019.

30. Christian Soize and Roger Ghanem. Probabilistic learning on manifolds constrained by nonlinear partial differential equations for small datasets. Computer Methods in Applied Mechanics and Engineering, 380:113777, 2021.

31. Christian Soize and Roger Ghanem. Probabilistic learning on manifolds (plom) with partition. International Journal for Numerical Methods in Engineering, 123(1):268–290, 2022.

32. Dimitris G. Giovanis, Nikolaos Evangelou, Ioannis G. Kevrekidis, and Roger G. Ghanem. Enabling probabilistic learning on manifolds through double diffusion maps. 551:114663. ISSN 0021-9991. doi: 10.1016/j.jcp.2026.114663.

33. Christian Apitz, Gary D. Webb, and Andrew N. Redington. Tetralogy of fallot. 374(9699): 1462–1471. ISSN 0140-6736, 1474-547X. doi: 10.1016/S0140-6736(09)60657-7.

34. K. H. W. J. ten Tusscher and A. V. Panfilov. Alternans and spiral breakup in a human ventricular tissue model. American Journal of Physiology-Heart and Circulatory Physiology, 291(3):H1088–H1100, September 2006. ISSN 0363-6135. doi: 10.1152/ajpheart.00109.2006. Publisher: American Physiological Society.

35. Oliver J. Britton, Alfonso Bueno-Orovio, Karel Van Ammel, Hua Rong Lu, Rob Towart, David J. Gallacher, and Blanca Rodriguez. Experimentally calibrated population of models predicts and explains intersubject variability in cardiac cellular electrophysiology. 110 (23):E2098–E2105. doi: 10.1073/pnas.1304382110.

36. Raffaele Coppini, Cecilia Ferrantini, Lina Yao, Peidong Fan, Martina Del Lungo, Francesca Stillitano, Laura Sartiani, Benedetta Tosi, Silvia Suffredini, Chiara Tesi, Magdi Yacoub, Iacopo Olivotto, Luiz Belardinelli, Corrado Poggesi, Elisabetta Cerbai, and Alessandro Mugelli. Late sodium current inhibition reverses electromechanical dysfunction in human hypertrophic cardiomyopathy. 127(5):575–584. doi: 10.1161/CIRCULATIONAHA.112.134932. Publisher: American Heart Association.

37. Julie K. Shade, Adityo Prakosa, Dan M. Popescu, Rebecca Yu, David R. Okada, Jonathan Chrispin, and Natalia A. Trayanova. Predicting risk of sudden cardiac death in patients with cardiac sarcoidosis using multimodality imaging and personalized heart modeling in a multivariable classifier. Science Advances, 7:8020–8048, 7 2021. ISSN 23752548. doi: 10.1126/SCIADV.ABI8020/SUPPL_FILE/SCIADV.ABI8020_SM.PDF.

38. Carmeline J Dsilva, Ronen Talmon, Ronald R Coifman, and Ioannis G Kevrekidis. Parsimonious representation of nonlinear dynamical systems through manifold learning: A chemotaxis case study. Applied and Computational Harmonic Analysis, 44(3):759–773, 2018.

39. RR Coifman and S Lafon. Geometric harmonics: a novel tool for multiscale out-of-sample extension of empirical functions. Applied and Computational Harmonic Analysis, 21(1):31– 52, 2006.

40. Nikolaos Evangelou, Felix Dietrich, Eliodoro Chiavazzo, Daniel Lehmberg, Marina Meila, and Ioannis G Kevrekidis. Double diffusion maps and their latent harmonics for scientific computations in latent space. Journal of Computational Physics, 485:112072, 2023.

